# Investigation on NTC (No-Template Control) Amplification in real time PCR test of Boar DNA sample

**DOI:** 10.1101/2025.08.07.669038

**Authors:** Muhammad Roy Asrori, Gracela Noveline Christiana Telaumbanua, Siti Khoirunnisa, Surjani Wonorahardjo, Suharti Suharti

## Abstract

Amplification in the No-Template Control (NTC) is a critical issue in real-time PCR (qPCR) as it can compromise the validity of results. This study was conducted to investigate the source of an observed signal in the NTC of a qPCR assay designed for boar DNA detection. Methodologies included melt curve analysis to characterize the amplification product and standard curve analysis to assess assay performance. The results from the melt curve analysis definitively confirmed that the product amplified in the NTC was not a primer-dimer, indicating that the signal did not originate from non-specific primer interactions. Furthermore, the primer set clearly demonstrated efficient amplification of the target DNA sequence in positive samples. The overall assay performance was validated by a standard curve, which showed acceptable linearity with a coefficient of determination (R^2^) of 1. Collectively, these findings suggest that the qPCR assay is robust and specific. The amplification observed in the NTC is attributed to minute DNA contamination rather than methodological flaws like primer-dimer formation, confirming the high sensitivity of the developed test.

## Introduction

Real-time Polymerase Chain Reaction (qPCR) is a powerful and indispensable technology in molecular biology, enabling the sensitive detection and precise quantification of specific nucleic acid sequences. Its capacity for rapid analysis, high specificity, and quantitative accuracy has made it a foundational method in diverse fields, from clinical diagnostics and genetic research to environmental monitoring and forensic science (Bustin, 2004). A particularly critical application of qPCR lies in food authenticity and safety testing, where it is employed for species identification. In this domain, qPCR assays for detecting porcine (boar or pig) DNA are paramount for verifying product claims, especially for food products intended for halal and kosher markets, where the absence of pork is a strict religious and regulatory requirement (Ali et al., 2012).

To ensure the validity and reliability of qPCR data, a stringent quality control framework is essential. The No-Template Control (NTC) is a cornerstone of this framework. The NTC reaction contains all the components of a standard qPCR—master mix, primers, and probes—but deliberately excludes the DNA template.4 Its primary function is to serve as a sentinel for contamination.5 In a properly executed experiment, the NTC should produce no amplification signal, thereby confirming that the reagents, consumables, and laboratory environment are free from contaminating DNA that could lead to false-positive results. The inclusion of NTCs is a non-negotiable standard stipulated by best practice guidelines, such as the Minimum Information for Publication of Quantitative Real-Time PCR Experiments (MIQE) (Bustin et al., 2009).

A significant and recurring challenge in qPCR is the appearance of an amplification signal in the NTC, a phenomenon known as NTC amplification. This undesirable outcome immediately calls the integrity of the entire experiment into question. It indicates the presence of contaminating nucleic acids, which can originate from a variety of sources:

✓ Reagent Contamination: Commercially available reagents, including the DNA polymerase or master mix, can sometimes contain trace amounts of bacterial or other environmental DNA (Borst et al., 2004).
✓ Carryover Contamination: The most common source is often DNA from previous experiments, particularly highly concentrated positive samples or amplicons (the products of previous PCRs), which can become aerosolized and contaminate pipettes, bench surfaces, and reagents (Aslanzadeh, 2004).
✓ Operator or Environmental Contamination: DNA from laboratory personnel or the surrounding environment can be inadvertently introduced during sample preparation or reaction setup.

Given the exponential nature of PCR amplification, even a single molecule of contaminant DNA can be amplified to detectable levels, resulting in a false-positive signal in the NTC.

This investigation specifically addresses the persistent issue of NTC amplification within a real-time PCR assay designed for the detection of boar DNA. The stakes in this particular application are exceptionally high. A false-positive result, potentially caused by NTC contamination, could lead to the incorrect rejection of a food product, resulting in significant economic losses and damage to brand reputation. The high sensitivity required to detect trace amounts of porcine material makes these assays particularly susceptible to contamination issues. Therefore, understanding and eliminating the root causes of NTC amplification is not merely a technical refinement but a critical necessity for ensuring the accuracy, reliability, and defensibility of the test results.

This study aims to systematically investigate the potential sources of contamination that lead to NTC amplification in a boar DNA-specific qPCR assay. By meticulously examining reagents, workflows, and environmental factors, we will identify the origins of the contaminating DNA and evaluate practical, effective strategies for its mitigation and prevention. The ultimate goal is to establish a robust protocol that eliminates NTC amplification, thereby safeguarding the integrity of boar DNA detection and reinforcing the trustworthiness of this vital analytical method. ⍰

## Materials and Methods

The materials for this study included nuclease-free water (NFW), forward and reverse primers specific for the target sequence, a commercial qPCR master mix (SimplyGreen qPCR Master Mix Kit Cat. No. SQ201-0020), genomic DNA from a boar sample (positive control), and genomic DNA from a cow sample (negative control). The primary instruments used in this study were: a set of calibrated micropipettes, s UV sterilization cabinet (UV Cleaner, Biosan), a microcentrifuge (Eppendorf 5427R), a vortex mixer (heidolph), a mini-centrifuge (for spin-downs), a real-time PCR thermal cycler(Quanstudio3, applied biosystem).

### Sample preparation

Sample preparation was performed in a UV cabinet. All materials were homogenized by vortexing and brief centrifugation. The reaction mixture was prepared according to the components detailed in Table 1.

**Table 1.** Mixture composition of sample preparation

| Sample | Volume of Boar DNA (μL) | Volume of Primer F (μL) | Volume of Primer R (μL) | Volume of NFW (μL) | Volume of Mastermix (μL) |
| --- | --- | --- | --- | --- | --- |
| Boar | 8 | 0,5 | 0,5 | 1 | 10 |
| Boar (10<sup>-1</sup> x) | 8 | 0,5 | 0,5 | 1 | 10 |
| Boar (10<sup>-2</sup> x) | 8 | 0,5 | 0,5 | 1 | 10 |
| Cow | 8 | 0,5 | 0,5 | 1 | 10 |
| NTC (NFW) | 0 | 0,5 | 0,5 | 9 | 10 |
| Without Primer F | 8 | 0 | 1 | 1 | 10 |
| Without Primer R | 8 | 1 | 0,5 | 1 | 10 |
| Without primer R-F | 8 | 0 | 0 | 2 | 10 |
| Without NFW | 9 | 0,5 | 0,5 | 0 | 10 |

### Real time q-PCR running

The real-time PCR (qPCR) was performed with the following thermal cycling protocol: an initial enzyme activation step at 95°C for 3 minutes (1 cycle), followed by 40 cycles of denaturation at 95°C for 15 seconds and a combined annealing/extension step at 60°C for 1 minute. Following the amplification stage, a melt curve analysis was conducted using the instrument’s default settings to verify the specificity of the amplified product (Bustin et al., 2009). Data acquisition and subsequent analysis were carried out using QuantStudio™ Design & Analysis Software v.1.5.3 (Thermo Fisher Scientific, 2021)

## Results and Discussion

### Amplification plot result of Boar DNA

The amplification results for the boar DNA real-time PCR (qPCR) assay are presented in Figure 1. The positive control containing boar DNA amplified robustly, with a quantification cycle (Cq) value of 18.47. However, late amplification was also unexpectedly observed in both the No-Template Control (NTC, Nuclease-Free Water) and the negative control (cow DNA), with Cq values of 25.97 and 26.18, respectively.

**Figure 1.**
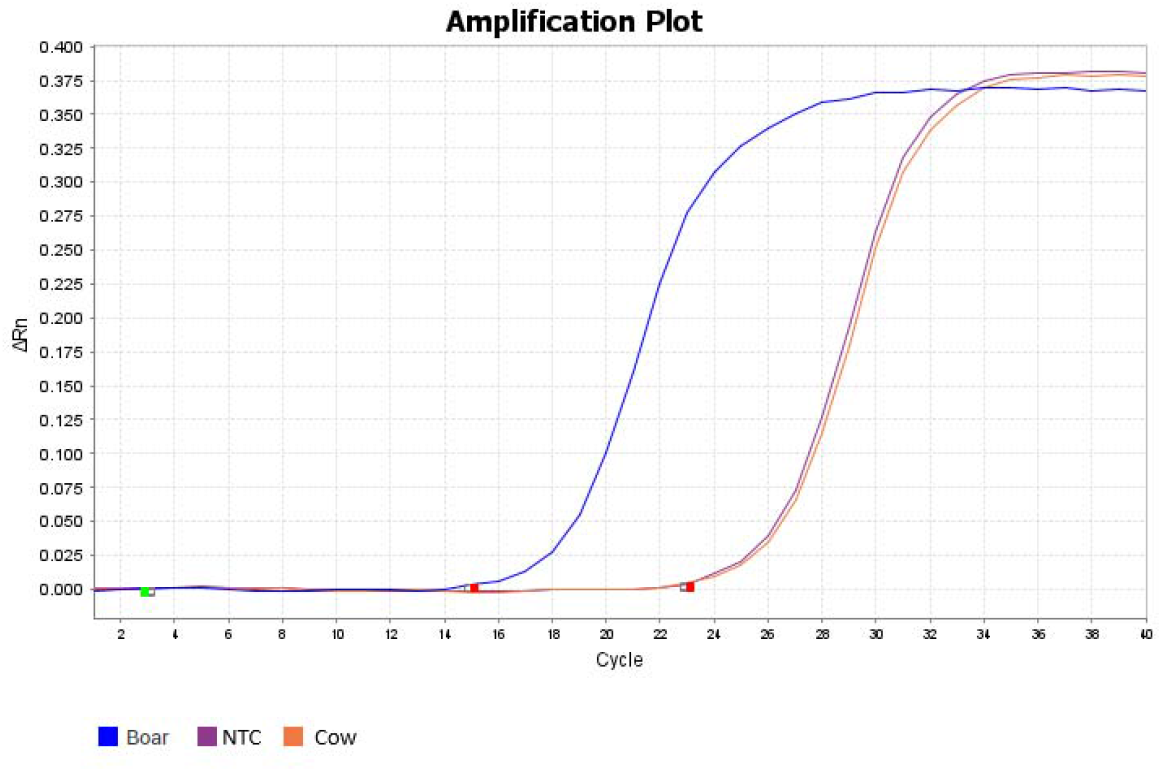
Amplification plot and Melt curve plot results of Boar DNA.

The presence of a signal in the NTC and the negative control samples indicates a potential issue with either reagent contamination or non-specific primer activity, requiring further investigation to ensure assay validity (Bustin et al., 2009). To diagnose the source of this amplification, a melt curve analysis was performed (Figure 2). The analysis revealed a single, distinct melt peak for all reactions, including those from the NTC and cow DNA samples. This result strongly suggests that the amplification was not caused by the formation of primer-dimers, which would typically produce a separate peak at a lower melting temperature (Ririe et al., 1997). Therefore, the late amplification observed in the control samples is most likely attributable to low-level contamination with the target DNA sequence rather than a flaw in primer design.

**Figure 2.**
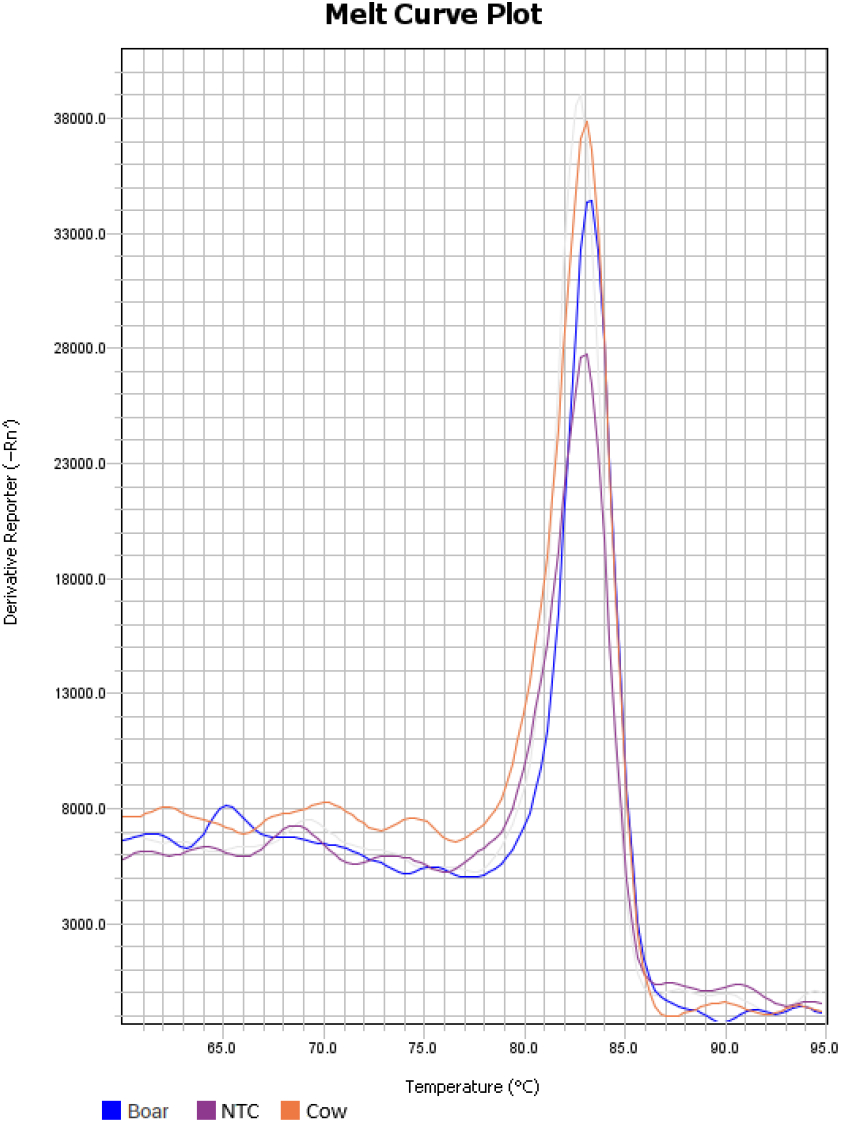
Melt curve plot of Boar DNA.

### Probing the Existence of the Primers

To validate that amplification was dependent on the complete primer set, a series of control reactions were conducted, with the results displayed in Figure 3. As shown, no amplification signal was detected in the control reactions that lacked both primers (no-primer control), contained only the forward primer, or contained only the reverse primer.

**Figure 3.**
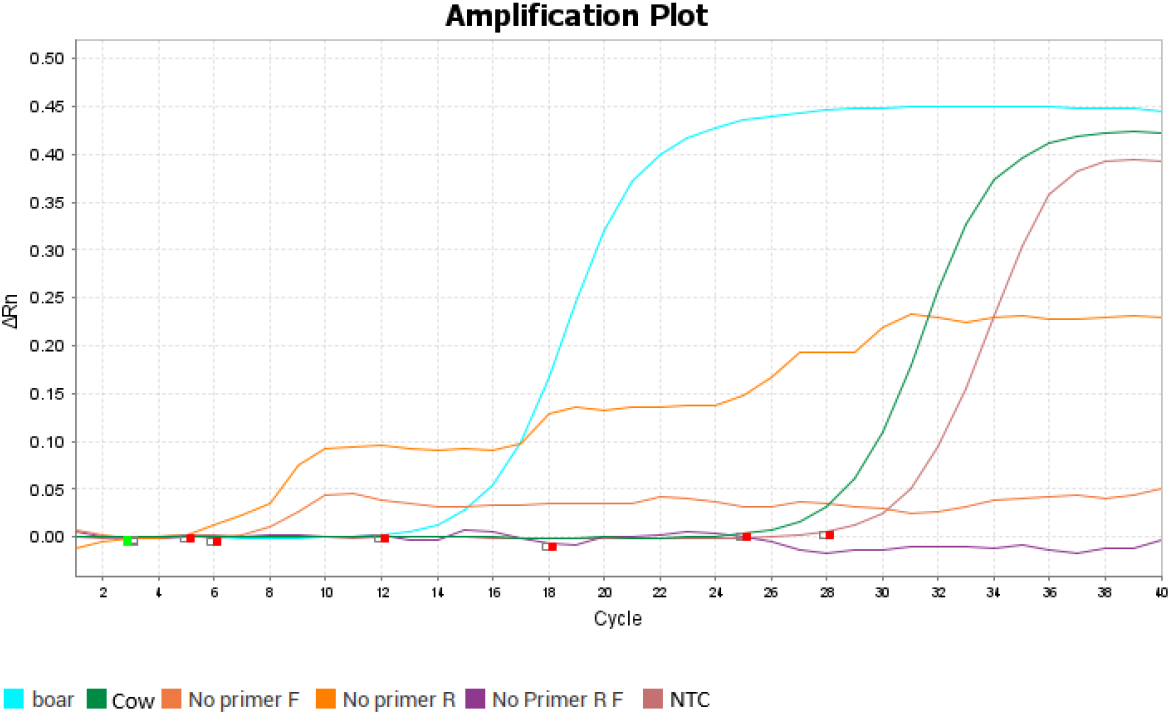
Amplification plot for evaluation of the primers in this study.

Successful amplification occurred exclusively in the reaction containing the complete primer pair (both forward and reverse). This outcome confirms that both the forward and reverse primers are essential for the reaction and that they function together to specifically amplify the target sequence. This is a fundamental confirmation of assay specificity and a critical validation step in PCR protocol development (Sambrook & Russell, 2001; Bustin et al., 2009)

Figure 4 displays the melt curve analysis conducted after amplification to assess product specificity. The results show that the positive control (boar), the negative control (cow), and the No-Template Control (NTC) all exhibited a single, dominant peak at the same melting temperature (Tm).

**Figure 4.**
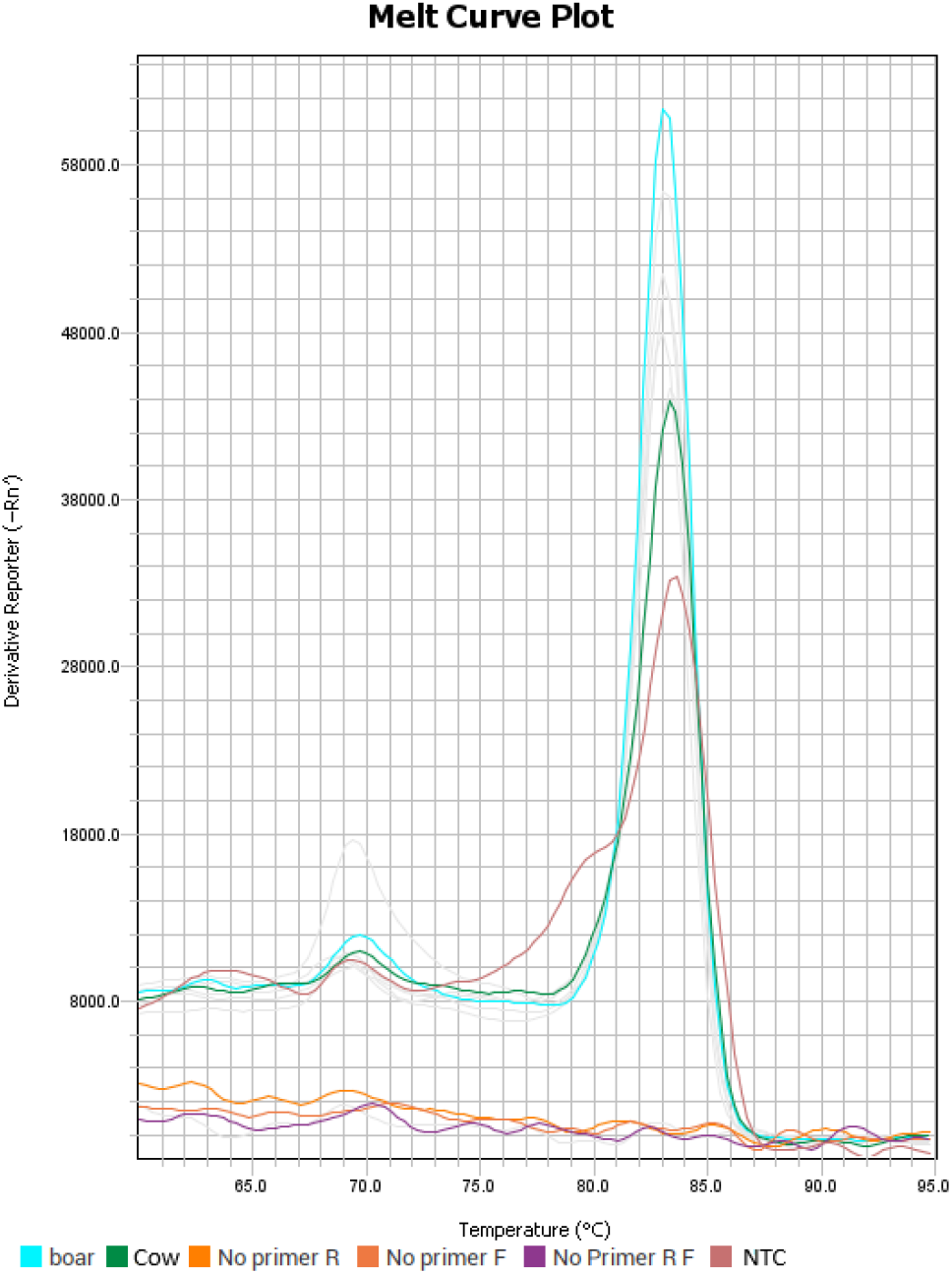
Melt curve plot for evaluation of the primers in this study.

This observation is critical. The presence of a single peak confirms that a single, specific DNA product was amplified in each reaction, effectively ruling out the formation of primer-dimers, which would typically appear as a distinct peak at a lower temperature (Ririe et al., 1997). More importantly, because the melt peaks from the negative control and NTC are identical to the peak from the positive boar sample, it confirms that the exact same DNA sequence was amplified in all reactions. This provides strong evidence that the unexpected amplification in the controls stems from low-level contamination with the target (boar) DNA, rather than from non-specific primer activity (Bustin et al., 2009).

### Standard curve of Boar DNA sample

To evaluate the amplification efficiency and linearity of the assay, a standard curve was generated using a serial dilution of boar DNA, with the results presented in Figures 5-7. The analysis yielded a linear equation of y = -2.064x + 21.5, where ‘y’ is the quantification cycle (Cq) and ‘x’ is the log of the initial DNA concentration.

**Figure 5.**
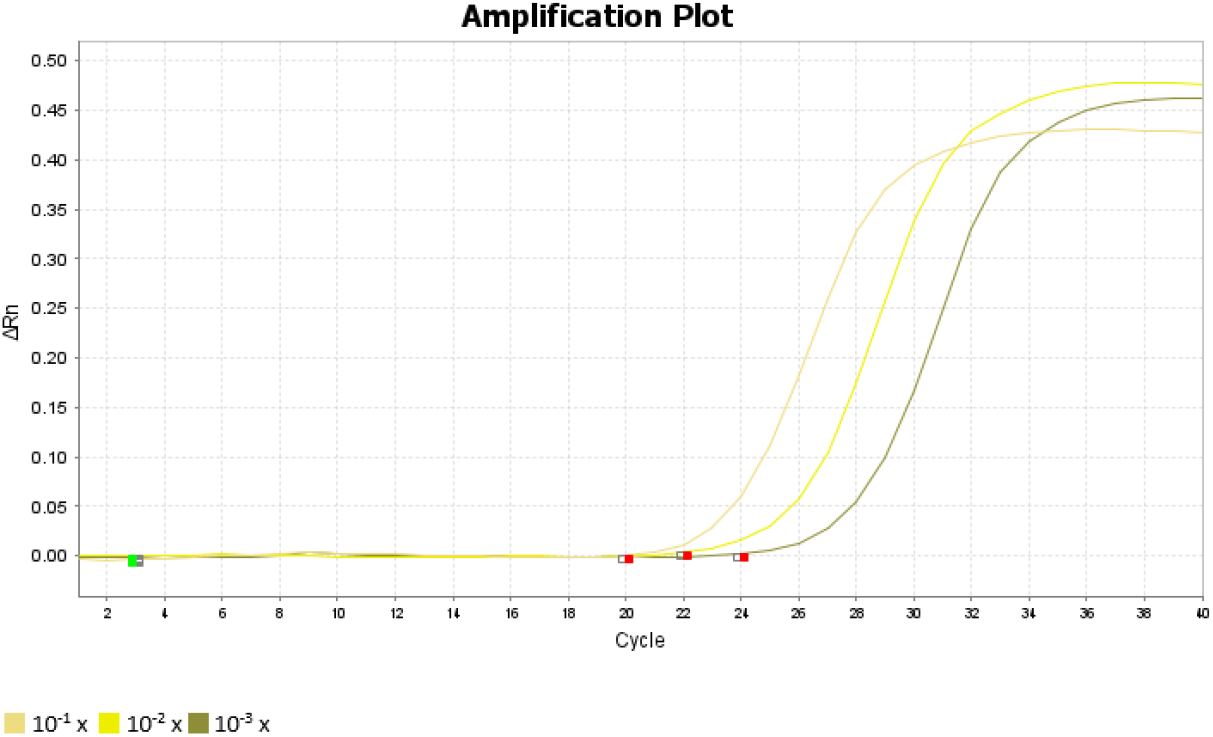
Amplification plot of the boar DNA variation.

**Figure 6.**
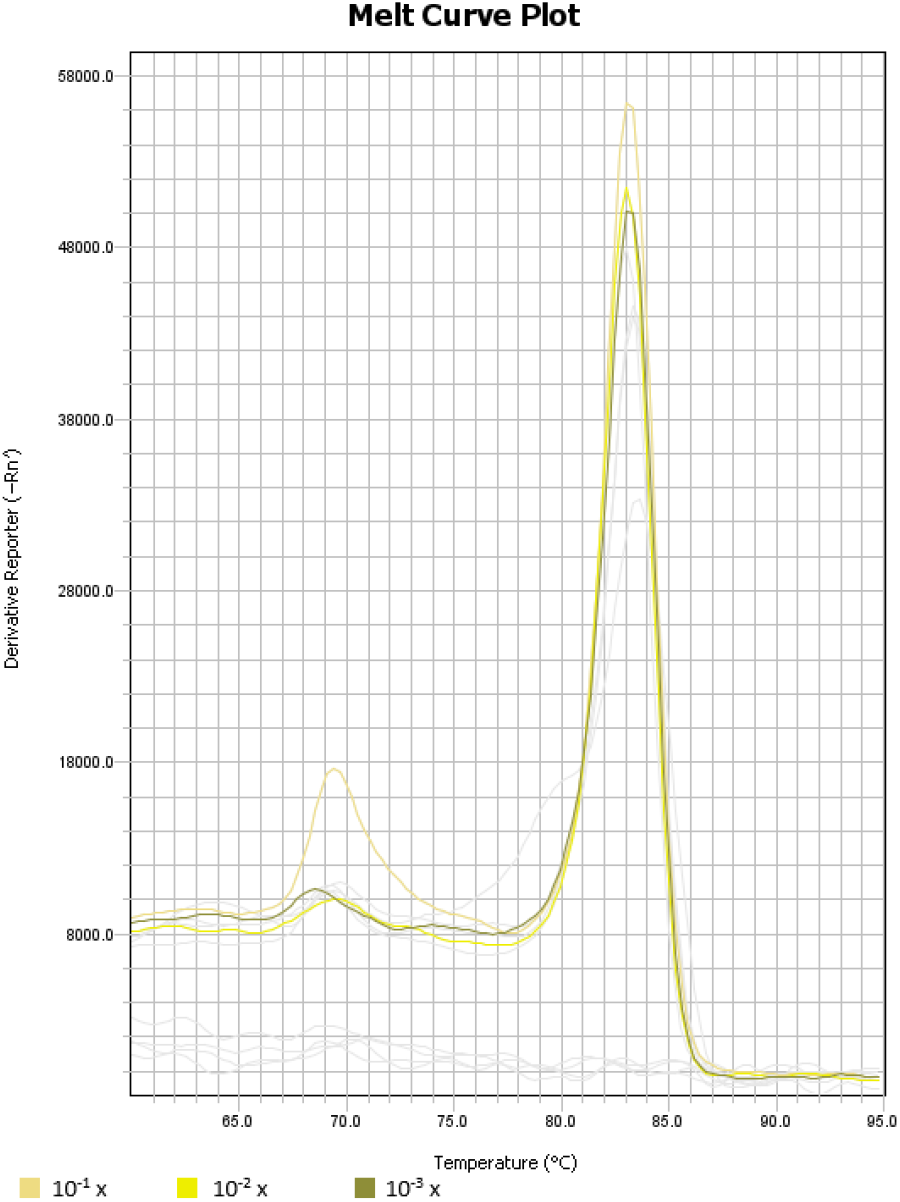
Melt curve plot of the Boar DNA variation.

**Figure 7.**
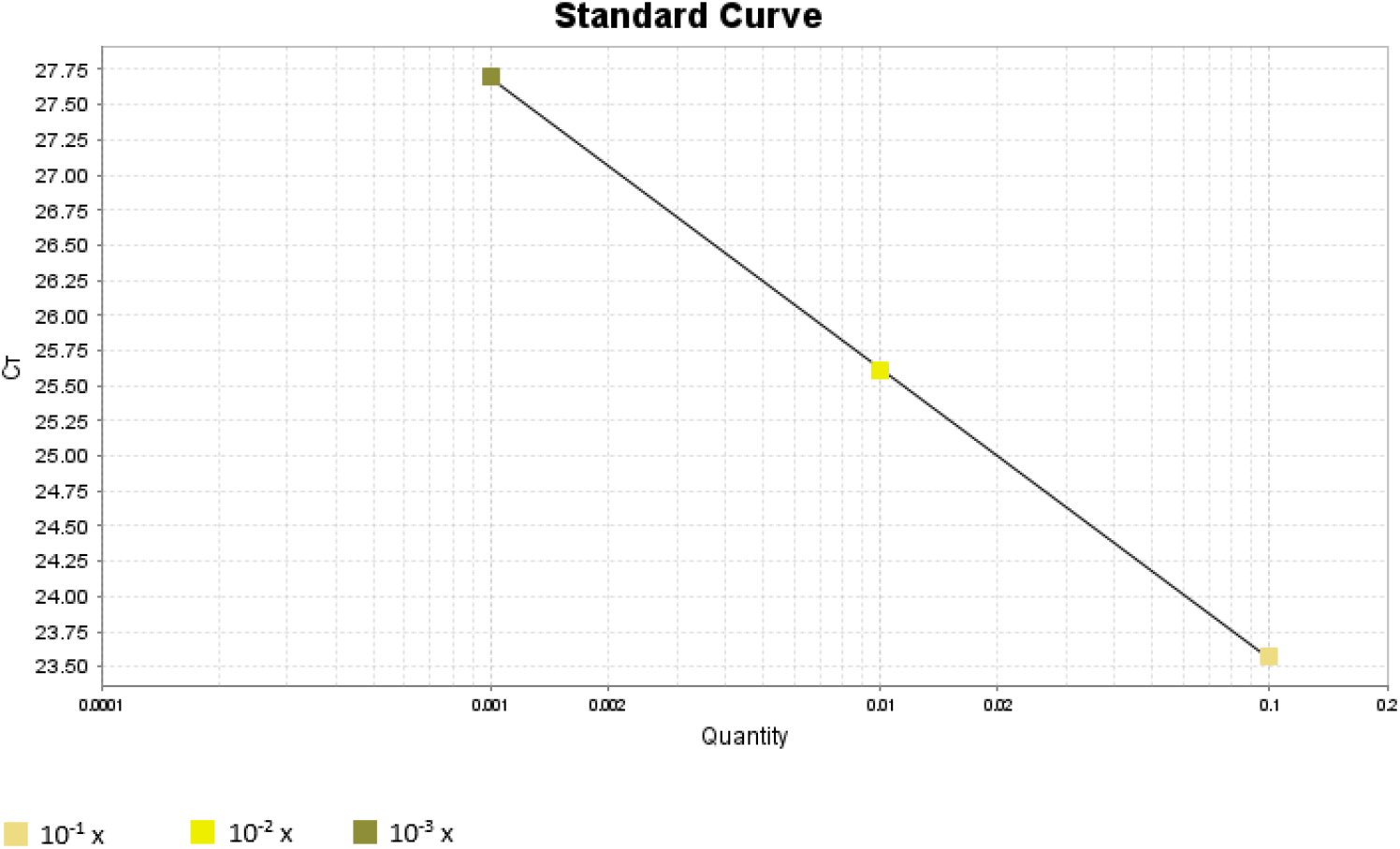
The Standard Curve results of the Boar DNA variation.

The key performance metrics derived from the curve were

1. Slope: -2.06
2. Coefficient of Determination (R2): 1.0
3. PCR Efficiency (Eff%): 205.15%

While the R^2^ value of 1.0 indicates perfect linearity across the tested concentration range, the slope and the resulting efficiency are outside of acceptable limits. An ideal slope for a qPCR assay is approximately -3.32, corresponding to 100% efficiency. An efficiency of 205.15% is significantly higher than the acceptable range of 90-110% and strongly suggests the presence of PCR inhibitors in the more concentrated standards or potential pipetting errors during the serial dilution (Bustin et al., 2009; Taylor et al., 2019). Therefore, despite the excellent linearity, the unreliable efficiency indicates that the assay requires further optimization before it can be used for accurate quantification.

## Conclusion

This investigation successfully diagnosed key issues within the real-time PCR assay for boar DNA. The analysis confirmed that the primers are functional and effectively amplify the target sequence. Melt curve analysis revealed that the signal observed in the No-Template Control (NTC) was not caused by primer-dimer formation, but rather by the amplification of a specific product, strongly indicating low-level DNA contamination. Furthermore, while the standard curve demonstrated perfect linearity (R2=1.0), the calculated amplification efficiency was unacceptably high (over 200%). This suggests the presence of PCR inhibitors in the concentrated samples or pipetting inaccuracies. In summary, the primary issues identified are reagent/environmental contamination and poor amplification efficiency, both of which must be resolved to ensure the assay is reliable for diagnostic use. The future work should focus on two main areas: contamination control and assay optimization.

## Acknowledgement

Thanks to Laboratory of Minerals and Advanced Materials that supports this study.

## Notes

### Competing Interest Statement

The authors have declared no competing interest.

